# A computational model for angular velocity integration in a locust heading circuit

**DOI:** 10.1101/2024.05.13.593806

**Authors:** Kathrin Pabst, Evripidis Gkanias, Barbara Webb, Uwe Homberg, Dominik Endres

## Abstract

Accurate navigation often requires the maintenance of a robust internal estimate of heading relative to external surroundings. We propose a novel model for angular velocity integration to update the representation of heading in the central complex of the desert locust. In contrast to similar models proposed for the fruit fly, this circuit model uses a single 360^*°*^ heading direction representation and is updated by neuromodulatory angular velocity inputs. Our computational model was implemented using steady-state firing rate neurons with dynamical synapses. The circuit connectivity was constrained by biological data and remaining degrees of freedom were optimised with a machine learning approach to yield physiologically plausible neuron activities. We demonstrate that the integration of heading and angular velocity in this circuit is robust to noise. The heading signal can be effectively used as input to an existing insect goal-directed steering circuit, adapted for outbound locomotion in a steady direction that resembles locust migration. Our study supports the possibility that similar computations for orientation may be implemented differently in the neural hardware of the fruit fly and the locust.

**Author summary:** In both fruit flies and locusts, a specific brain region has been observed to have an activity pattern that resembles a compass, with an activity peak moving across an array of neurons as the animal rotates through 360 degrees. However, some apparent differences in the properties of this pattern between the two species suggest there may be differences in how this internal compass is implemented. Here we focus on the locust brain, building a computational model that is based on observed neural connections and using machine learning to tune the system. Turning by the simulated locust provides modulatory input to the neural circuit that keeps activity in the array aligned to its heading direction. We simulate a migrating locust that tries to keep the same heading despite perturbances and show this circuit can steer it back on course. Our model differs from existing models of the fruit fly compass, showing how similar computations could have different implementations in different species.

Table 1.
Abbreviations for neuron types and brain regions in the desert locust (*Schistocerca gregaria*) and homologues in the fruit fly (*Drosophila melanogaster*).

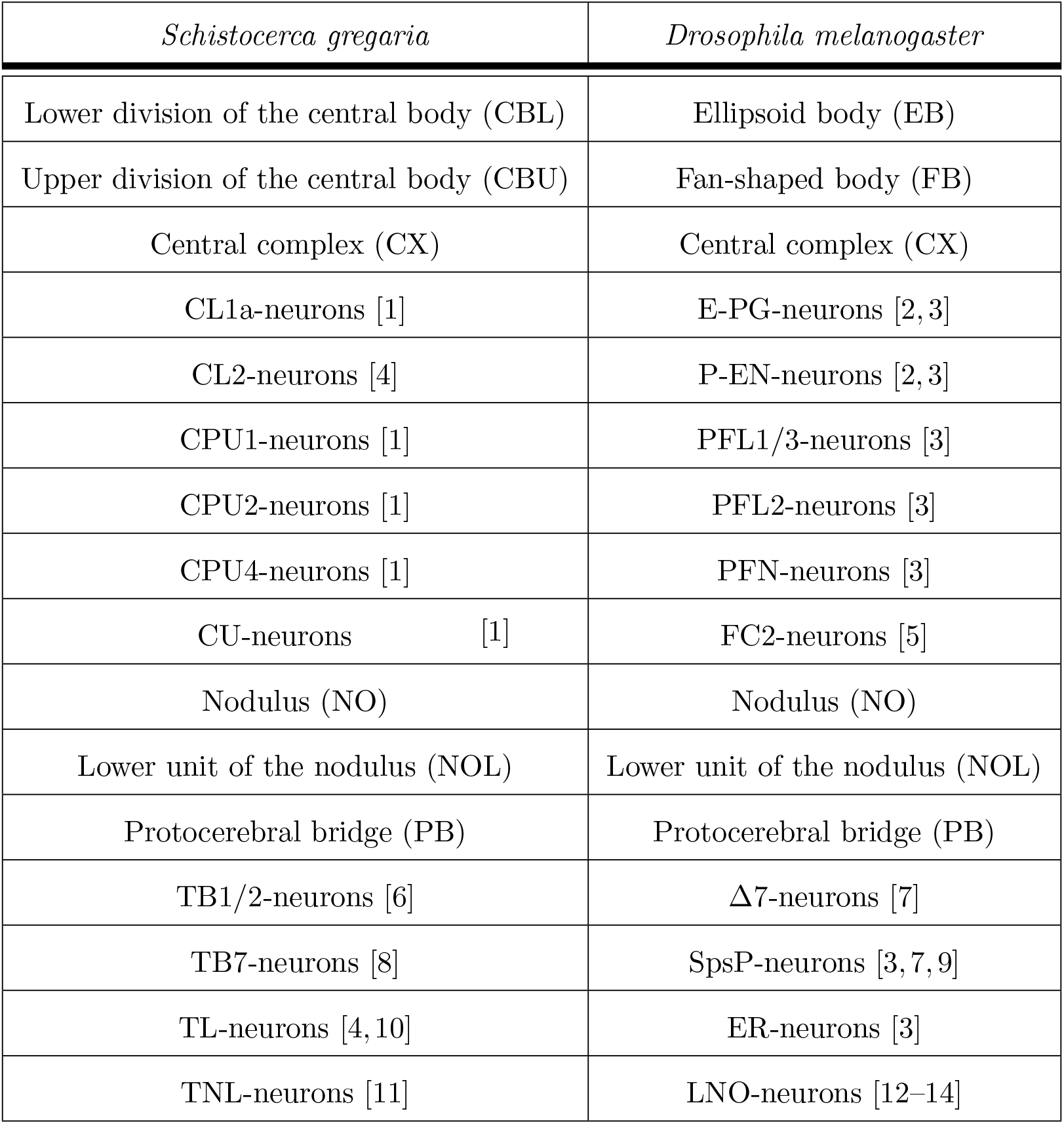

## Introduction

Various navigational strategies have evolved across diverse ecological contexts, many relying on a robust estimate of the animal’s current heading direction [15–17]. Shared across species (including humans) [18, 19], these strategies likely stem from similar neuronal and computational foundations. Investigating orientation and its neural substrates in a model organism provides a gateway to uncovering general mechanisms of spatial cognition. With their impressive navigational abilities and suitability for both laboratory and field studies, insects emerge as excellent model organisms for investigating navigation [20].

The navigation centre of the insect brain is located in the central complex (CX) [21–24]. This brain region is a midline-spanning group of four major neuropils: The protocerebral bridge (PB), the upper (CBU) and lower (CBL) divisions of the central body (also known as the ellipsoid body (EB) and fan-shaped body (FB) in some species), and the paired noduli (NO) [22]. The PB, CBU, and CBL are compartmentalised into columns and the CBU, CBL, and NO are stratified into layers [1, 21, 25]. These columns and layers are interconnected by tangential and columnar neurons following stereotypical projection patterns [1, 9]. Tangential neurons provide multimodal inputs from various brain regions to the CX [3, 8, 26–28], while columnar neurons connect columns between the different neuropils and serve as the principal output sites of the CX [11, 22]. This organization is highly conserved, and tight structure-function relationships reveal the biological implementation of vector-based algorithms in the CX [24].

Analogous to head direction cells [15] observed in mammals [29, 30], numerous CX-neurons respond to celestial cues and map the animal’s heading direction relative to the angle of polarised skylight and the solar azimuth [11, 15, 31]. To track the animal’s orientation robustly, these cells integrate partly redundant inputs from various modalities [23], including self-motion generated signals [2, 32] such as efference copies or optic flow. In the fruit fly, columnar E-PG-neurons form a comprehensive 360^*°*^ compass within the EB, with calcium imaging revealing a single activity maximum or compass bump encoding the animal’s heading direction [2, 32]. Projection schemes of E-PG-neurons yield one 360^*°*^ compass representation in each hemisphere of the PB, shifted relative to each other by 22.5^*°*^ [2, 3, 33].

The representation of space in the fruit fly CX exhibits variability between individuals and across contexts [31, 32]. In contrast, in the desert locust *Schistocerca gregaria*, intracellular recordings suggest a single 360^*°*^ compass encoding across the entire width of the PB [6, 34, 35], and projection patterns of columnar neurons imply two intercalated 180^*°*^ representations of space along the CBL [1, 36]. The data suggest that the compass topography is consistent across individual locusts. Figure 1A illustrates the internal compass topographies in the PB of the fruit fly and the desert locust. In addition to this functional difference between the fruit fly and locust heading circuits, many structural differences exist [37, 38]. A major difference is the ring-shaped EB in the fruit fly, which is a striking exception to the crescent-shaped homologous neuropils in most other insect species [39], such as the CBL in the locust. Given the prevalence of this locust-like CBL architecture, we deemed it relevant to investigate the consequences of a locust-like 1 *×* 360^*°*^ heading representation in the PB. Many models of insect navigation [37, 40–42] assume a 2 *×* 360^*°*^ heading representation across the two halves of the PB, but this pattern might be an exception in the fruit fly rather than common to all insects.

**Fig 1.**
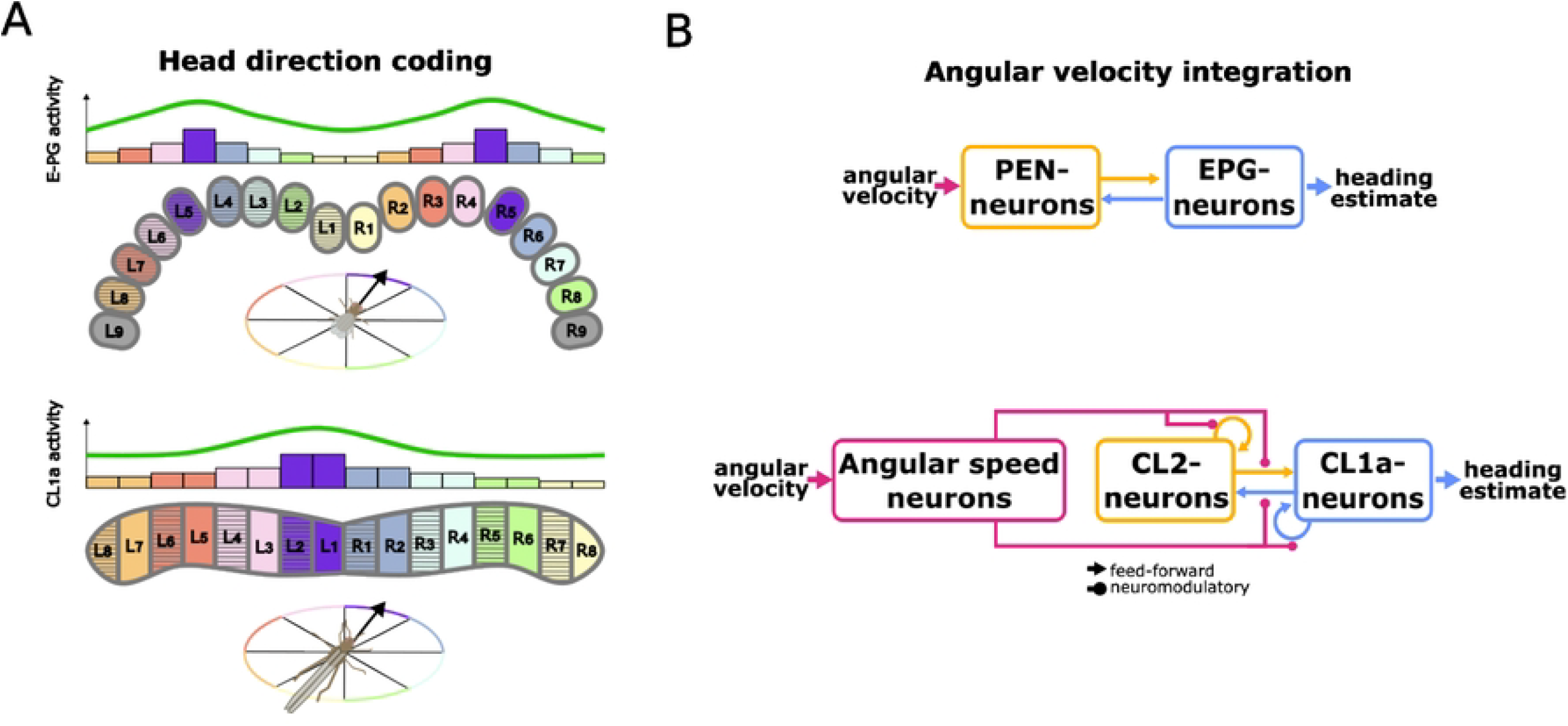
Overview of the proposed model. (A) Schematic comparison of heading encoding in the fruit fly and the desert locust protocerebral bridge (PB), based on data from [13, 33] and [36] and inspired by illustrations from [3, 24, 35]. Columnar E-PG- and CL1a-neurons encode the heading direction (indicated by colour and arrow orientation) of the insect. Bar graphs illustrate the activity level of neurons in each PB column, revealing a sinusoidal pattern of activity across the PB. In contrast to the fruit fly’s 2 *×* 360^*°*^ representation of space with two activity maxima (compass bumps) along the PB, one on either side, our model of the locust heading circuit assumes a 360^*°*^ spatial map with a single compass bump along the entire PB. (B) Diagrammatic comparison of information flow through the fruit fly heading circuit proposed by [2] and our proposed model of the desert locust heading circuit. Both circuits feature homologous columnar neurons (E-PG- and P-EN-neurons / CL1a- and CL2-neurons). In the fruit fly heading circuit proposed by [2], P-EN activity directly depends on the animal’s angular velocity. In our proposed locust heading circuit, an abstract class of angular speed neurons modulates the circuit connectivity depending on the animal’s angular velocity.

A model of angular velocity integration in the fruit fly heading circuit [2] includes E-PG-neurons encoding heading and P-EN-neurons conjunctively encoding heading and angular velocity. Within this model, asymmetric activation of P-EN-neurons in the two halves of the PB occurs based on the fruit fly’s turning direction, resulting in a shift of E-PG and P-EN activity maxima in the EB and the PB through the circuit’s connectivity. Homologous columnar neurons in the locust, CL1a- and CL2-neurons, display projection schemes [1] suggesting similar connectivity, although excitation and inhibition remain uncertain. Functionally, CL1a-neurons encode heading relative to a fixed point of reference [34, 35], and recordings from CL2-neurons suggest directional sensitivity to rotational optic flow [36]. The circuit likely receives angular velocity inputs from tangential neurons [11].

In previous work, using a simplified model with linear neural units, discrete-time updates and binary rotation encoding (left vs. right) we showed that the observed heading encoding in the locust could in principle be shifted appropriately by a multiplicative rotation-dependent modulation of the firing rate [36, 43]. Here, we significantly extend this work by developing a firing rate model of the locust heading circuit with synaptic dynamics and optimising its function under structural and biologically plausible parameter constraints. An overview of the included neuron types and their interactions is shown in Figure 1B. We were interested in determining whether such a constrained model would be able to integrate a continuum of angular velocities and generate locust-like neural activity and orientation behaviour. This approach is conceptually related to the ‘bounded rationality’ models in cognitive science [44] where realistic behaviour emerges by training models constrained by available resources towards optimal behaviour. The results show that a heading circuit with a different compass topography from the fruit fly, and with neuromodulatory instead of feed-forward angular velocity inputs, can still function to maintain a robust heading estimate and to control steering behaviour.

## Model and methods

The heading circuit model and all simulations were implemented in Python 3.11. We used the machine learning library PyTorch (version 2.2.1) for optimisation of the model’s free parameters as described in section ‘Free parameters and their optimisation’ .

### Neuron model

The network consists of single-compartment steady-state firing rate neurons abstracted from integrate-and-fire neurons [45]. Since the main focus of our work is the circuit topology, we only present the key assumptions regarding free and constrained parameters here. For a complete derivation of the neuron model, please refer to the supplementary material section ‘Neuron model derivation’.

The dynamics of the model neurons are governed by two time constants: *τ*_*m*_, the membrane time constant, and *τ*_*s*_, the synaptic time constant. Estimates for *τ*_*m*_ vary with neuron type. While *τ*_*m*_ = 1.5 ms is common in cortical neurons [45], the membrane time constants used in the fruit fly CX model of [2] are larger by an order of magnitude to capture observed delays between E-PG and P-EN activity in walking flies [2]. See supporting information section ‘Functional transmission delays between CL1a- and CL2-neuron populations’ for an order-of-magnitude delay estimation approach. We chose to model these delays by slow synapses, i.e. *τ*_*s*_ *≫ τ*_*m*_. This assumption justifies the use of a steady-state firing rate model with an explicit dynamics model for the synapses. The steady-state potential *U*_*∞*_ of the membrane is

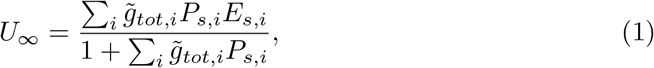

where *P*_*s,i*_ is the post-synaptic ion channel opening probability, *E*_*s,i*_ is the synaptic reversal potential, and 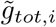 is the total relative synaptic conductance of synapse *i*. This conductance is calculated as

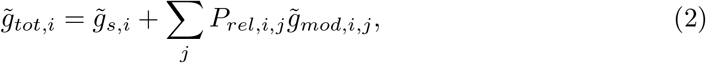

where 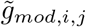 and 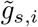 are its two contributions. Note that 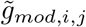 is multiplicatively modulated by pre-synaptic transmitter release probability *P*_*rel,i,j*_ from modulatory input *j*, and 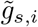 is not modulated. Both contributions are free parameters, hereafter referred to as ‘weights’, that will be optimized subject to connectivity constraints. We compute the firing rate *r* with a logistic sigmoid activation function

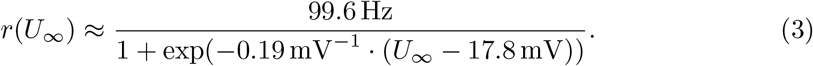

We model the effect of a spike on *P*_*s,i*_ by a single-exponential kernel. The kernel is convolved with the density of the pre-synaptic spike train, which we assume to have inhomogeneous Poisson process statistics with rate *r*_*i*_. We argue that the Poisson assumption is approximately valid for CX neurons, since *r*_*i*_ does not exceed 50 Hz in available data from the locust [36]. This implies that the typical inter-spike interval is substantially longer than the refractory period. Following the derivation in [45], the time course of *P*_*s,i*_ can then be described by the differential equation

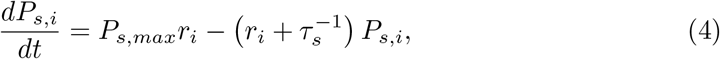

where *P*_*s,max*_ is the maximum synaptic open probability, which we set to 1.

The circuit model relies on multiplicative neuromodulation for heading representation updates. Neurons with a modulatory effect change their pre-synaptic transmitter release probability *P*_*rel,i,j*_ proportionally to their rate and with time constant *τ*_*s,mod*_. We expect *τ*_*s,mod*_ *> τ*_*s*_ because neuromodulation often involves signal transmission cascades.

### Heading circuit model

The heading circuit consists of columnar CL1a- and CL2-neurons and an abstract class of angular speed neurons. The latter are sensitive to angular velocity and modulate the connectivity among CL1a- and CL2-neurons.

The CX receives diverse inputs from tangential neurons, supposedly including explicit angular speed information [11]. As there are no data indicating which specific tangential neurons assume this role in the locust, we included abstract, functionally inspired angular speed neurons in our model. These units are designed to summarise the characteristics of biologically identified tangential neurons, potentially delivering inputs to the PB, CBL, and the lower unit of the NO (NOL). We modelled one angular speed neuron tuned to clockwise rotation (AS_cw_) and one tuned to counterclockwise rotation (AS_ccw_). Their firing rates are given by

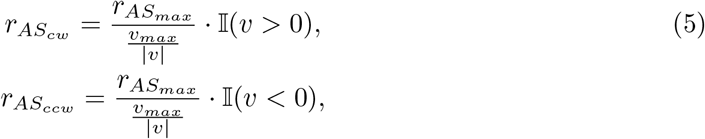

where 𝕀(.) is the indicator function which is 1 if the argument is ‘true’ and 0 otherwise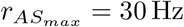 is the maximum firing rate of angular speed neurons, which corresponds to a high firing rate of CX-neurons in the locust under experimental conditions [36].

Based on data from flying locusts responding to striped patterns moving at up to 90 ^*°*^*/*s [46], we conservatively assume *v*_*max*_ = 150 ^*°*^/s is the maximum angular velocity of a locust.

CL1a-neurons encode the animal’s orientation in a compass-like manner [6, 34, 35]. The preferred heading directions of the 16 CL1a-neurons included in our model are 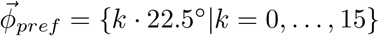. Here, k corresponds to the PB column index a neuron arborises in. PB columns are indexed from left to right, i.e., L8 has index 0 and R8 has index 15 (cf. Figure 1A for labelling of PB columns). The distribution of preferred heading directions of CL1a-neurons along the PB is based on the distribution of preferred solar azimuths derived from sky polarisation tuning along the PB in four types of CX neurons (CL1-, TB1-, CPU1-, and CPU2-neurons in locusts—E-PG-, *Δ*7-, PFL1/3-, and PFL2-neurons in fruit flies) [35]. This results in a 360^*°*^ representation of space with a single activity maximum or compass bump along the PB (cf. Figure 1A, lower panel, for a schematic representation). The firing rates of CL1a-neurons are initialised with a sinusoidal relationship to the initial heading *ϕ*(*t*_0_) at time *t*_0_:

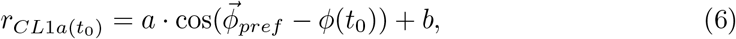

where the rate amplitude *a* = 5 Hz and the operating point *b* = 25 Hz were determined from the data reported in [36].

CL2-neurons inherit heading information from CL1a-neurons. In addition, they are sensitive to rotational optic flow compatible with yaw rotation [36]. CL2-neurons and homologous neurons in the fruit fly, P-EN-neurons [2, 33], show opposite directional selectivity with neurons in the left PB hemisphere preferring counterclockwise rotations and neurons in the right PB hemisphere preferring clockwise rotations. In our model, angular velocity and direction information enter through the angular speed neurons. They modulate the weights of synapses from CL1a-onto CL2-neurons and of synapses within the subset of CL2-neurons such that firing rates of CL2-neurons change in an angular velocity-dependent manner.

To maintain a stable operating point b (see Equation 6), all CL1a- and CL2-neurons receive an additional input from a bias neuron constantly firing at ca. 100 Hz.

#### Neuronal projections and connectivity assumptions

The heading circuit connectivity was derived from anatomical projection data, based on two assumptions: first, smooth fibre endings indicate input regions of CX-neurons and varicose fibre endings indicate output regions; second, overlapping arbors with opposite polarity are potentially synaptically connected. See Figures 2A-B for the general projection schemes of the modelled neuron types. The connectivity implications that follow from these two assumptions are detailed below, together with the respective evidence.

**Fig 2.**
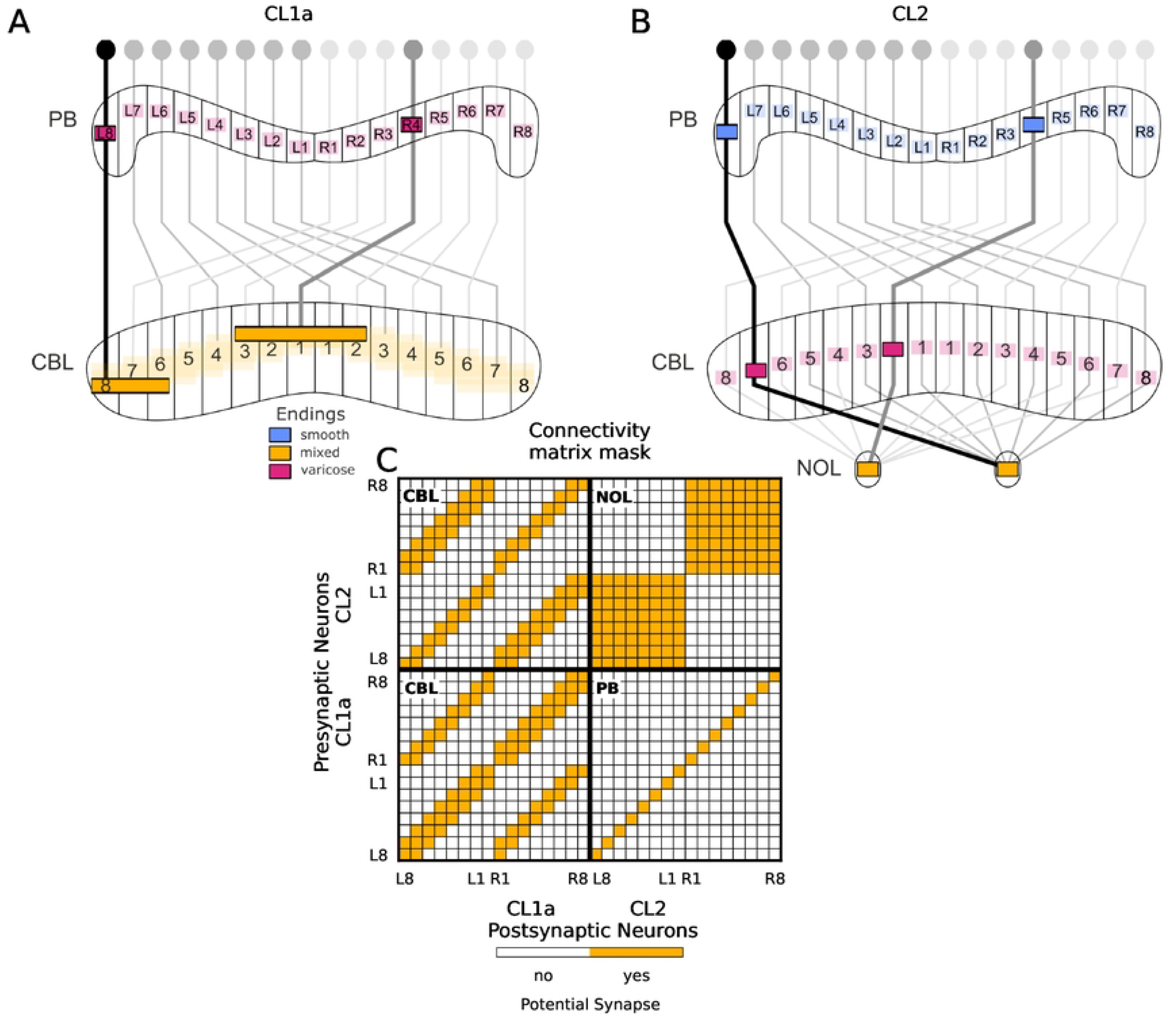
Deduction of the heading circuit connectivity from anatomically plausible connections in the CX. (A,B) Projection schemes of CL1a- and CL2-neurons, adapted from [1]. CL1a-neurons connect multiple adjacent columns of the CBL to single columns of the PB. CL2-neurons connect single columns of the PB to single columns of the CBL and to the contralateral NOL. The pattern by which PB and CBL are connected is shifted by one column when comparing the two neuron types. (C) Connectivity matrix mask for CL1a- and CL2-neurons, indicating potential synapses (yellow squares) without specifying excitation versus inhibition or strength. Neurons are arranged according to their position in the PB. PB, protocerebral bridge; CBL and CBU, lower and upper division of the central body, NOL, lower unit of the noduli.

First, CL1a-neurons provide input to CL2-neurons in the PB. Each CL1a-neuron has varicose endings in a single PB column, and each CL2-neuron has smooth endings in a single PB column [1], such that each CL1a-neuron could provide input to the CL2-neuron arborising in the same column of the PB. In the fruit fly, E-PG-neurons also provide input to P-EN-neurons in the PB [2].

Second, CL2-neurons provide input to CL1a-neurons in the CBL. Each CL2-neuron has varicose endings in a single CBL column. Each CL1a-neuron has central bleb-like endings in a single CBL column surrounded by smooth endings in up to two columns on both sides of it (‘mixed’ endings in Figures 2A) [1]. Both CL1a- and CL2-neurons connect columns of the PB to columns of the CBL, and both neuron types project columns in each half of the PB onto alternating columns across the entire width of the CBL. The projection schemes of the two neuron types are shifted by one column (cf. the pattern of alternating colours across the columns of the CBL in Figure 2A,B). Each CL2-neuron could thus provide input to CL1a-neurons arborising in the ipsilateral as well as in the contralateral half of the PB but in the same column of the CBL. In the fruit fly, P-EN-neurons provide input to E-PG-neurons in the EB [2]. However, neuron projections differ significantly between the two species, with each E-PG-neuron innervating one of 16 wedges of the EB (corresponding to one column of the CBL) and each P-EN-neuron innervating one of eight tiles of the EB (corresponding to two neighbouring columns of the CBL).

Third, neighbouring CL1a-neurons make synaptic contacts in the CBL. The organization of varicose terminals in a single column of the CBL flanked by smooth endings in neighbouring columns renders it likely that CL1a-neurons in adjacent PB columns are synaptically connected in the CBL. Due to their projection schemes detailed above, each CL1a-neuron could provide input to other CL1a-neurons arborising in both the ipsilateral and the contralateral half of the PB. Synaptic contacts between E-PG-neurons have also been demonstrated in the EB of the fruit fly [3, 47].

Fourth, CL2-neurons arborising in the same NOL provide input to each other. In addition to the PB and the CBL, CL2-neurons arborise in the contralateral NOL. In the fruit fly, all P-EN neurons from one hemisphere of the PB are connected with each other in the NO [3, 47], and we assume the same is true for CL2-neurons from one hemisphere in the NOL.

Lastly, angular speed neurons potentially modulate all synapses among CL1a- and CL2-neurons. Tangential neurons provide inputs to the CX from various other brain regions [8], and many types of tangential neurons have been immunostained for neuromodulatory transmitters. Specifically, TB6-/TB7-neurons innervating the PB have been immunostained for tyramine and the neuropeptides orcokinin and locustatachykinin [48–50]. TB8-neurons (also innervating the PB) have been immunostained for octopamine [49]. TL1- and certain TL4-neurons innervating the CBL have been immunostained for the neuropeptide orcokinin [50], and TN-neurons innervating the NO have been immunostained for tyramine [49]. In our model, we summarised these properties in angular speed model-neurons, enabling them to up- and down-regulate synapses among columnar neurons.

Figure 2C shows the resulting connectivity matrix mask for all potential synapses among CL1a- and CL2-neurons in the model. There are no data indicating the inhibitory or excitatory nature of these proposed synapses. Since data on their strength is not available either, we determined all synaptic weights using an optimisation algorithm (see section ‘Free parameters and their optimisation’).

#### Stimuli

We supplied two types of input stimuli to the neural circuit; heading and angular velocity. The heading stimulus was provided only once, to initialise network activity at the beginning of each experiment. The initial firing rates of the ensemble of CL1a-neurons as well as the ensemble of CL2-neurons were set to encode a particular heading direction via Equation 6. This choice of identical CL1a- and CL2-activities was motivated by the observation that P-EN and E-PG activity maxima align if the angular velocity is very low [2]. Angular velocity inputs were provided to the angular speed neurons throughout simulations and were used to continuously update the heading signal encoded in the CL1a-neuron population activity. The angular speed neurons initially fired at a rate corresponding to an angular velocity *v*(*t*_0_) = 0 ^*°*^/s. The initial states of all synapses in the network were set to a stationary state that is reached when the circuit receives zero angular velocity inputs for a long time (i.e. much longer than *τ*_*m*_ and *τ*_*s*_).

### Free parameters and their optimisation

The free parameters of the model were the synaptic weights (cf. equation 1 and equation 2) of all potential synapses (indicated by the connectivity mask, see Figure 2C), including both the feed-forward synapses between CL1a- and CL2-neurons and the modulatory effects of angular speed neurons on these synapses. These parameters were determined via unconstrained gradient-based optimisation with the L-BFGS [51] algorithm. Gradients were computed with PyTorch’s automatic differentiation algorithm. All synaptic weights were optimised so that the network reproduced activity targets encoding the true heading during or after an integration time interval of 200 ms angular velocity integration interval. We used a 4^th^ order Runge-Kutta integrator [52] to integrate the system of ordinary differential equations. Integration time steps from 1 to 8 ms yielded comparable results.

For a random initial heading direction *ϕ*(*t*_0_), we computed the true heading at time *t*_*n*_, *ϕ*(*t*_*n*_), by integrating the angular velocity *v*(*t*_*n*_):

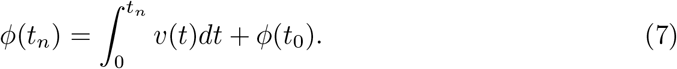

To train the network to maintain a stable heading encoding when *v*(*t*) = 0 ^*°*^*/*s (const.) throughout the integration time interval, we generated one maintenance activity target 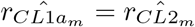 per random heading direction. The optimisation algorithm then minimised the mean squared error between this target and the network output at every 10th integration time steps. Simultaneously, to train the network to shift the heading representation when receiving nonzero angular velocity inputs, we applied one of 128 randomly drawn constant *v*(*t*) *∈* [*−*150 ^*°*^/s, 150 ^*°*^/s] for 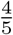 of the integration time interval, and *v*(*t*) = 0 ^*°*^/s for the remaining 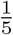th of the integration time interval afterwards. The optimisation algorithm then minimised the mean squared error between the shift activity targets 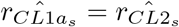 resulting from integrating the angular velocity in the final 10 integration time steps and the network outputs. This choice of identical CL1a- and CL2-targets was again motivated by the observation that P-EN and E-PG activity maxima align if the angular velocity is very low [2].

Since there is no unique solution to this optimisation problem, we regularised the minimum with a low-entropy prior. The regulariser promoted similar values for all synaptic weights across repeated connectivity structures in the PB and the CBL, i.e., it punished variance along the diagonals of the quadrants of the connnectivity matrix. This regularization was intended to produce a visually pleasing appearance of the connectivity matrix [53]. The relative weight of the regulariser was 0.1.

In Python pseudo-code, the complete objective function used to optimise the synaptic weights is

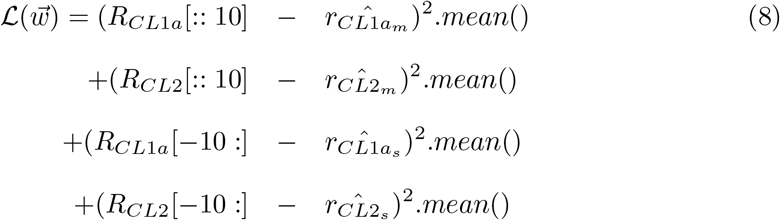

where 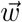 is the vector of all free parameters, or weights (cf. equation 2) and *R*_*CL*1*a*_ is the matrix of the rates of the 16 CL1a-neurons at each time predicted by the model. *R*_*CL*1*a*_[:: 10] indicates the rates of all CL1a-neurons at every tenth integration time step, and *R*_*CL*1*a*_[*−*10 :] indicates the rates at the final ten integration time steps. 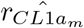 is the vector of target rates during heading maintenance and 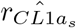 is the vector of target rates after shifting the heading direction with angular speed input and likewise for the CL2 rates.

### Simulations and evaluation

#### Evaluating the noise robustness of heading and angular velocity integration

We first explored the effects of altering membrane potentials, synaptic release probabilities, and synaptic weights on the accuracy of the integration of heading and angular speed. We varied the membrane potentials by adding zero-mean Gaussian noise with standard deviation *σ*_*U*_ *∈ {*0.0, 0.1, 0.5, 1.0*}*mV at every millisecond. We sampled Beta-distributed noise for the synaptic release probabilities with the mean equal to the noise-free value, and a pseudocount *∈ {*10, 100, 1000*}*(These pseudocounts correspond to a coefficient of variation approximately *∈ {*0.1, 0.01, 0.001*}*). We chose a Beta distribution because it is range-limited to [0, 1], which is important for an interpretation as a probability. Finally, we randomly perturbed synaptic weights by uniformly distributed multiplicative noise with range *∈ {*0.01, 0.03, 0.05*}* at the start of each integration trial, effectively applying noise to both modulatory and non-modulatory synaptic weights relative to the original weight strength. We carried out N = 2000 integration trials lasting 4000 ms for each noise value. In each trial, the circuit activity was initialised based on a random heading direction via Equation 6. The angular speed neurons received angular velocity inputs generated from lowpass filtered Gaussian noise, to mimic the observed trajectories of walking locusts. We quantified the accuracy of the heading circuit via the average angular error between the true heading (cf. Equation 7) and the heading estimate encoded in the activity of CL1a-neurons:

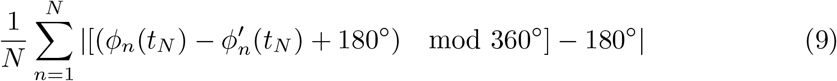

where *ϕ*(*t*_*N*_) and *ϕ*^*′*^(*t*_*N*_) are the ground truth and estimated heading directions at the final points in time of each trial. Estimated heading directions *ϕ*^*′*^(*t*_*N*_) were computed as the phase of the closest fitting cosine to CL1a-neuron activity. The fit was obtained by a linear regression of the cosine values to the rates *r*_*CL*1*a*_(*t*_*N*_) with arbitrary amplitude and baseline scaling.

#### Evaluating the attractor stability of the compass states

We further explored the ability of the circuit to converge to a stable compass-like heading encoding after a perturbation of presynaptic rates and synaptic weights. We added Gaussian noise with a standard deviation relative to the amplitude of 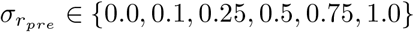 to presynaptic input to the network at the beginning of the simulation. We also randomly perturbed synaptic weights by uniformly distributed multiplicative noise with range *∈ {*0.05, 0.1, 0.15*}* at the start of each integration trial. We carried out N = 1000 integration trials lasting 100 ms for each value. In each trial, the circuit activity was initialised via Equation 6 based on a random heading direction. Throughout the trial, angular speed neurons received angular velocity inputs generated from lowpass filtered Gaussian noise. We quantified the stability of the heading circuit via the mean-squared deviation between the CL1a-neurons’ activity and the best-fitting sine at the end of each trial.

#### Testing of the heading circuit in an agent simulation

Lastly, we tested whether the heading circuit could guide locomotor behaviour. To simulate a walking locust in a simulated world, we linked the heading circuit to a circuit that produces outputs for goal-directed steering [41]. In short, the heading circuit outlined above updates an internal heading representation by integrating angular velocity information (cf. Figure 3A). We adapted the steering circuit to produce steering signals by comparing representations of the current heading and a constant goal direction (cf. Figure 3B). In each trial, we initialised the heading circuit’s activity based on a heading direction at time (t_0_). At all subsequent points in time t_1:N_, the updated heading representation served as an input to the steering circuit which drove a simulated motor system that moved the agent. The agent’s behaviour resulted in angular velocities that were fed back into the heading circuit (cf. Figure 3C). To connect the heading circuit to the steering circuit, we had to make several adjustments to the original model of [41]: First, the CL1a output of the heading circuit was transformed via a logistic sigmoid to match the rounded square wave of CL1 activities of the goal-directed steering circuit. Second, in the original steering circuit, the current heading direction is represented by two compass bumps across 16 CL1a-neurons. This is transformed into a single bump across 8 TB1-neurons. We optimised the CL1a-TB1 connectivity in the steering circuit, to achieve the same with the single bump heading representation in the 16 CL1a-neurons of our model. The assumptions and optimization targets are detailed in the following: the optimization was constrained by projection data from CL1a-[1] and TB1-neurons in the locust [8, 54] (cf. Figure 3D). As outlined above, we assumed that smooth and varicose fibre endings indicate input- and output sites and that overlapping fibres with opposite polarity indicate potential synapses.

**Fig 3.**
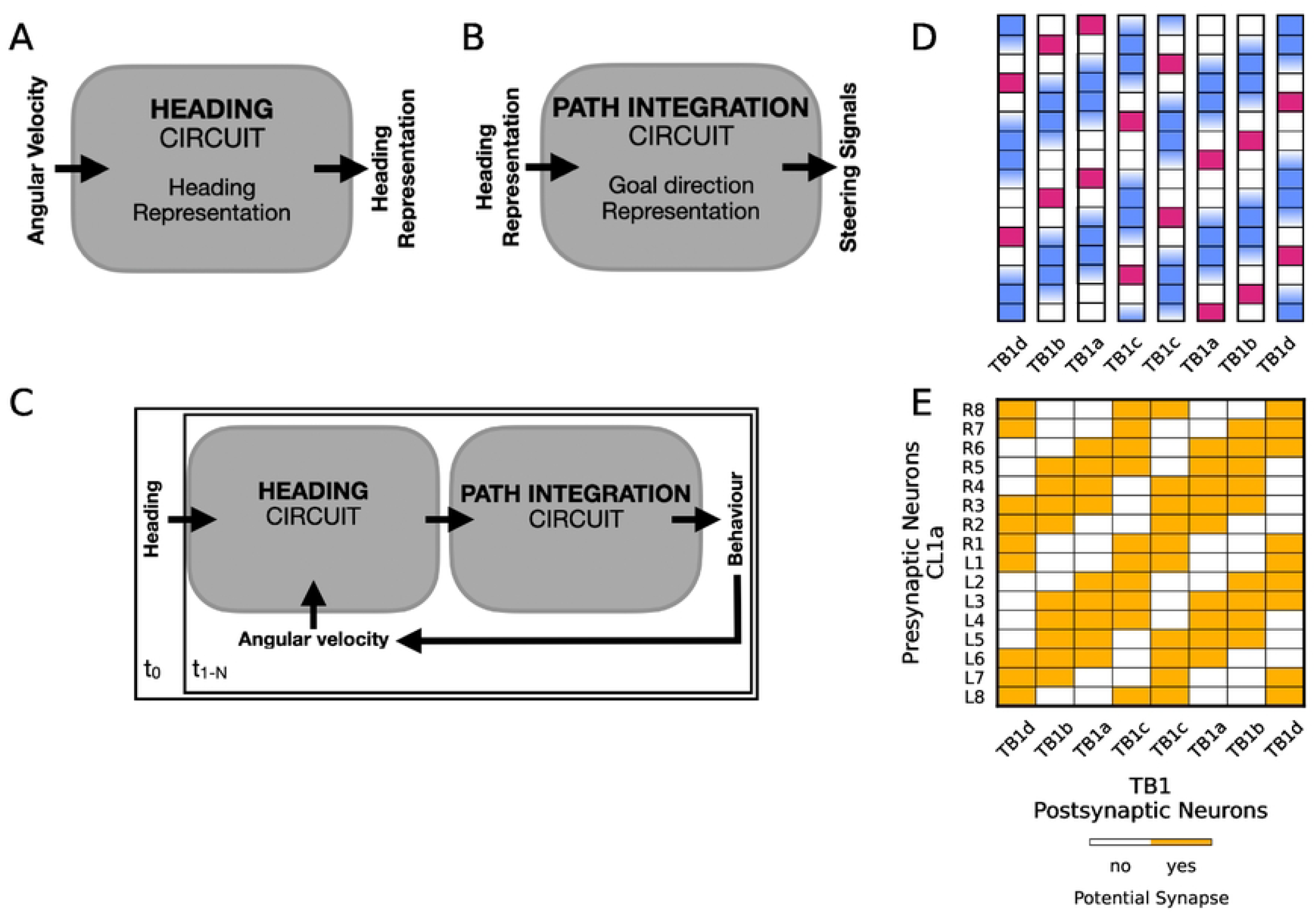
Connecting the heading circuit to the goal-directed steering circuit. (A) The heading circuit introduced above updates an internal heading representation by integrating angular velocity inputs. (B) The goal-directed steering circuit proposed by [41] produces steering signals to align the current heading with a goal direction representation. (C) The two circuits can be linked to form a closed loop. Starting with an initial heading representation (at time t_0_) the heading representation produced by the heading circuit is used as an input to the steering circuit, which is initialised with a constant goal direction. The behavioural output of the steering circuit in turn produces angular velocity input for the heading circuit (at all consequent times t_1:N_). (D) Topographic organization of TB1-neuron subtypes in the PB, adjusted from [54]. Magenta squares indicate varicose fibre endings, blue indicates smooth fibre endings. (E) Connectivity matrix mask for CL1a- and TB1-neurons within the steering circuit, adjusted for projection data from the locust illustrated in panel D (TB1-neurons) and Figure 2A (CL1a-neurons). Yellow squares indicate potential synapses without specifying excitation versus inhibition or strength. CL1a-neurons are arranged according to their position in the PB, TB1-neurons are labelled according to the subtypes with matching arborisations.

Figure 3E illustrates the resulting potential synapses from CL1a-onto TB1-neurons within the steering circuit. As there are no data indicating the weights of these potential synapses, we optimised the CL1a-TB1 connections to map rounded square wave activities across 16 CL1a neurons to similar activities across 8 TB1-neurons. The weights were constrained to be positive. Furthermore, we regularised the solution by a quadratic weight decay to push all unnecessary weights close to zero. Also, we implemented a -5^*°*^ phase shift between the CL1a and TB1 bumps, to compensate for biases introduced by rounding the continuous-time representation of our heading circuit to the discrete-time steering model. This implementation-dependent bias necessitated a slightly more liberal interpretation of the CL1a-TB1 connectivity scheme depicted in Figure 4D, see Figure 4E. However, we argue that this extended connectivity is still in agreement with the data of [54] within the error margins of that data.

**Fig 4.**
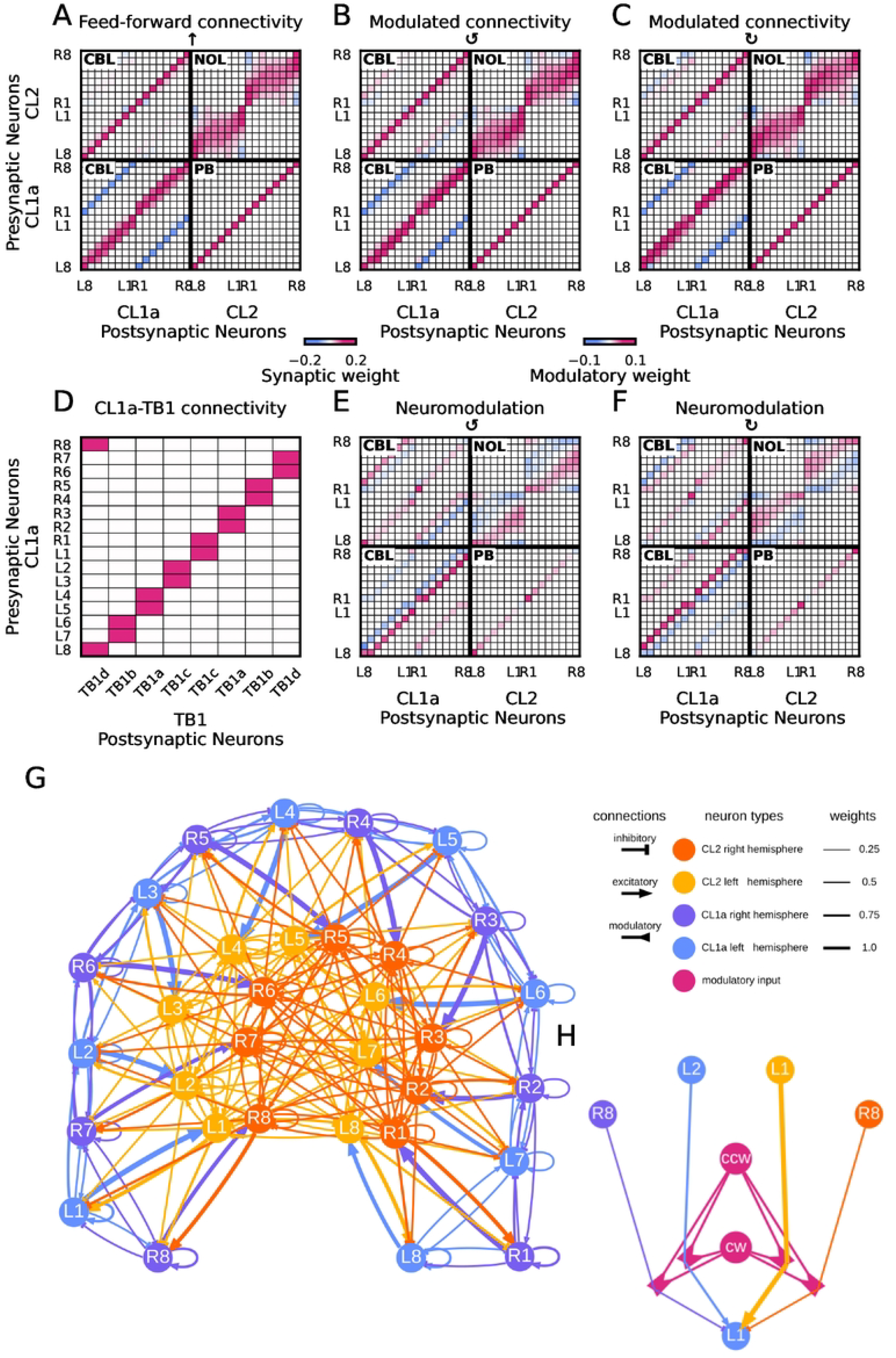
Biologically constrained and optimised feedforward connectivity and neuromodulation. (A,B,C) Effective connectivity of CL1a- and CL2-neurons. Neurons are arranged according to their position in the PB. Effective positive (excitatory) synaptic weights are displayed in magenta, negative (inhibitory) ones in blue (with the value range of the left colorbar). Colours saturate at 25% of the maximal synaptic weight value for better visibility. Panel A displays the connectivity matrix optimised to maintain a heading signal at zero angular speed. Panels B and C illustrate the optimised, modulated circuit connectivity during counterclockwise and clockwise rotation, respectively. (E,F) Modulatory weights of angular speed neurons onto synapses between CL1a- and CL2-neurons during counterclockwise and clockwise rotation, respectively. Positive (upregulating) modulatory weights are displayed in magenta, negative (downregulating) ones in blue (with the value range of the right colorbar). Note that B and C are the sums of A+E and A+F, respectively. (D) Optimised connectivity of CL1a- and TB1-neurons within the steering circuit. CL1a-neurons are arranged according to their position in the PB, and the eight TB1-neurons are labelled according to the subtypes with matching arborisations. Excitatory synaptic weights are displayed in magenta (with the value range of the left colorbar). (G) Force-directed graph of the optimised effective connectivity for heading maintenance, matching the connectivity matrix shown in panel A. (H) Example microcircuit illustrating neuromodulation affecting CL1a-neuron L1. Recurrent self-connection not shown. The clockwise (cw) and counterclockwise (ccw) angular velocity inputs modulate the connections between CL1a- and CL2-neurons.

Third, we aimed to simulate outbound locomotor behaviour during the long-range phase of a long-distance navigational task [55]. During this phase, the animal maintains its goal direction based on global cues. In the goal-directed steering circuit, a homing vector is encoded in CPU4-neurons and updated continuously. The authors suggested that CPU4-neurons could encode a fixed direction during long-range migration in other insects, and we thus hard-coded a goal vector fixed in direction and length in this layer, instead of performing continuous updates.

The ability of the agent to maintain a steady travel direction was quantified by Equation 9, with *ϕ* and *ϕ*^*′*^ as the actual and the ideal heading direction (matching the goal direction) of the agent at the final point in time of the behavioural simulation. At the beginning of each trial, the agent was placed in the simulated world with a random heading direction. This initial heading was translated to the initial heading circuit activity r_CL1a_(t_0_) and r_CL2_(t_0_). A random and fixed goal direction was encoded in r_CPU4_(t_0:n_). The agent then performed 500 steps, with a total duration of 50 s. We explored the effect of displacements by wind on the agent’s performance. We modelled wind with two parameters: P(translation) is the probability of a gust of wind that displaces the agent laterally by the magnitude of one step with each step the agent takes. P(rotation) is the probability of a gust of wind that rotates the agent by a random magnitude with each step the agent takes. Each gust of wind lasted for a random duration in 500 to 1500 ms. Translation and rotation were mutually exclusive. We chose *P*(*translation*), *P*(*rotation*) *∈ {*0.0, 0.01, 0.02, 0.03, 0.04*}* and repeated N = 1000 runs for each value.

## Results

### A proposed circuit for heading and angular speed integration in the desert locust CX

The optimised circuit connectivity is illustrated in Figure 4. The CL1a-CL2 connectivity for maintaining a heading representation encoded in an activity pattern is displayed in Figure 4A. Colour saturation indicates unmodulated synaptic weight, i.e. 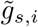 in equation 2. The solution features uniformly excitatory synapses from CL1a-onto CL2-neurons in the PB. In the CBL, CL1a-neurons excite CL1a-neurons arborising in the same PB arm, including excitatory self-connections, and inhibit CL1a-neurons projecting to the contralateral one. Also in the CBL, the same pattern emerges in synapses from CL2-onto CL1a-neurons, with additional inhibitory synapses between neurons arborising near the midline of the PB. In the NOL, CL2-neurons from the same hemisphere are interconnected with near excitation (including excitatory self-connections) and far inhibition.

Modulatory effects of angular speed neurons onto synapses between CL1a- and CL2-neurons induce a shift of the heading signal during turns. Figures 4B and C display the effective, modulated, connectivity (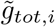 in equation 2) during counterclockwise and clockwise rotations. This connectivity results from adding modulatory weights 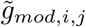 depicted in Figures 4E and F to the connectivity shown in Figure 4A, after scaling them with the pre-synaptic release probabilities *P*_*rel,i,j*_. The solution features neuromodulation of synapses between all neuron populations in the PB, in the CBL, and the NOL. The general pattern in this solution for counterclockwise turns is that, within the CBL, the synapses of CL1a-neurons onto other CL1a-neurons arborising in one column of the PB to the left become less excitatory, whereas synapses onto CL1a-neurons arborising in one column of the PB to the right become more excitatory. This modulation is reversed for clockwise turns. A similar pattern can be seen for the synapses between CL2-neurons within the NOL, except that the modulations extends across several columns of the PB. This solution requires angular speed neurons to exert both up- and down-regulating effects. Potential biological substrates for this dual effect are addressed in the discussion. The direction (up- or down-regulation) of modulatory effects is homogeneous across neuron populations (given by neuron type and hemisphere), but strengths vary. As for synapses from CL2-neurons onto CL1a-neurons and for synapses from CL1a-neurons onto CL2-neurons, modulation is more pronounced at synapses between neurons arborising in the outer- and innermost columns of the PB. A comparison of the effective connectivity at zero (Figure 4A) and nonzero angular velocity (see Figures 4B and C) reveals that modulation does not substantially alter overall excitation and inhibition in the circuit since modulatory effects are comparably weak.

The optimised CL1a-TB1 connectivity is shown in Figure 4D. Note that the connections far from the diagonal have been pruned, even though they would have been permissible (cf. Figure 3E). This means that only one of the two input domains of each TB1 neuron (cf. blue squares in Figure 3D) is functionally connected to a CL1a neuron. Whether this solution emerges from the noise-free CL1a activities used for the optimisation, the positivity constraint on the weights or the weak quadratic regulariser is a question for future investigations.

Figure 4G shows a force-directed graph of the feed-forward circuit connectivity shown in Figure 4A. Synaptic strength is indicated by edge thickness. Note that the synaptic strengths are up- and down-regulated during turns, but the polarity of synapses (excitatory or inhibitory) is not altered by neuromodulation. Figure 4H shows a microcircuit illustrating neuromodulation affecting CL1a-neuron L1, omitting the recurrent self-connection. The clockwise (cw) and counterclockwise (ccw) angular velocity inputs modulate the connections between columnar neurons (see Figures 4E,F).

The interplay of up- and down-regulation around the main diagonals of the connectivity matrix (in other words, the modulation of synapses between neurons in adjacent PB columns) serves a computational purpose, effectively yielding a discretised derivative of the sinusoidal CL1a activity pattern across the PB.

The derivative of a sine wave is a cosine wave (and vice versa). Any phase-shifted (co)sine wave can be computed by a weighted linear superposition of a sine and a cosine wave. This mathematical relationship is expressed by the trigonometric identity [56]:

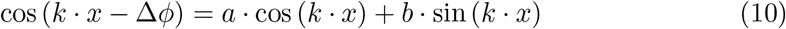

Here, *a* = cos(Δ*ϕ*), *b* = sin(Δ*ϕ*), 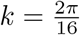 and *x ∈ {*0, …, 15*}* indexes PB columns from L8 to R8 (labelling as depicted in Figure 1A). Consequently, the connectivity modulation introduces a linear combination of a sine and a cosine wave, resulting in a shift of the compass bump.

### Accuracy and robustness of heading and angular velocity integration

The model was evaluated in three different simulations. We first assessed the capability of the heading circuit for updating an initial heading representation by integrating a time series of angular velocity inputs (see Figure 5A for an example). Throughout these simulations, both CL1a- and CL2-neurons consistently exhibited sinusoidal activity patterns, localising in a single maximum along the PB. The position of this compass bump aligned with the ground truth heading direction (Equation 7) and dynamically responded to angular velocity inputs. As in the model of the locust heading circuit proposed by [37], the compass bump demonstrated the ability to seamlessly transition between the lateral ends of the PB. To determine how much neuromodulation in the PB, CBL and/or NOL contribute to the accuracy of angular speed integration, we restricted modulatory inputs to either the CL1a -neurons or the CL2-neurons and re-optimized the network. We evaluated the average absolute integration error and its standard error at the simulation endpoint after 2000 trials of 4 s duration. Errors were 25.05^*°*^ *±* 0.39^*°*^ for the CL1a+CL2 modulated network, 26.83^*°*^ *±* 0.45^*°*^ for modulation of CL1a inputs only, and 26.91^*°*^ *±* 0.39^*°*^ for a network with modulation of the CL2 inputs only. These average errors indicate that neuromodulation of either or both neuron populations will allow for comparably accurate angular velocity integration, with a slightly higher accuracy for the CL1a+Cl2 modulated network.

**Fig 5.**
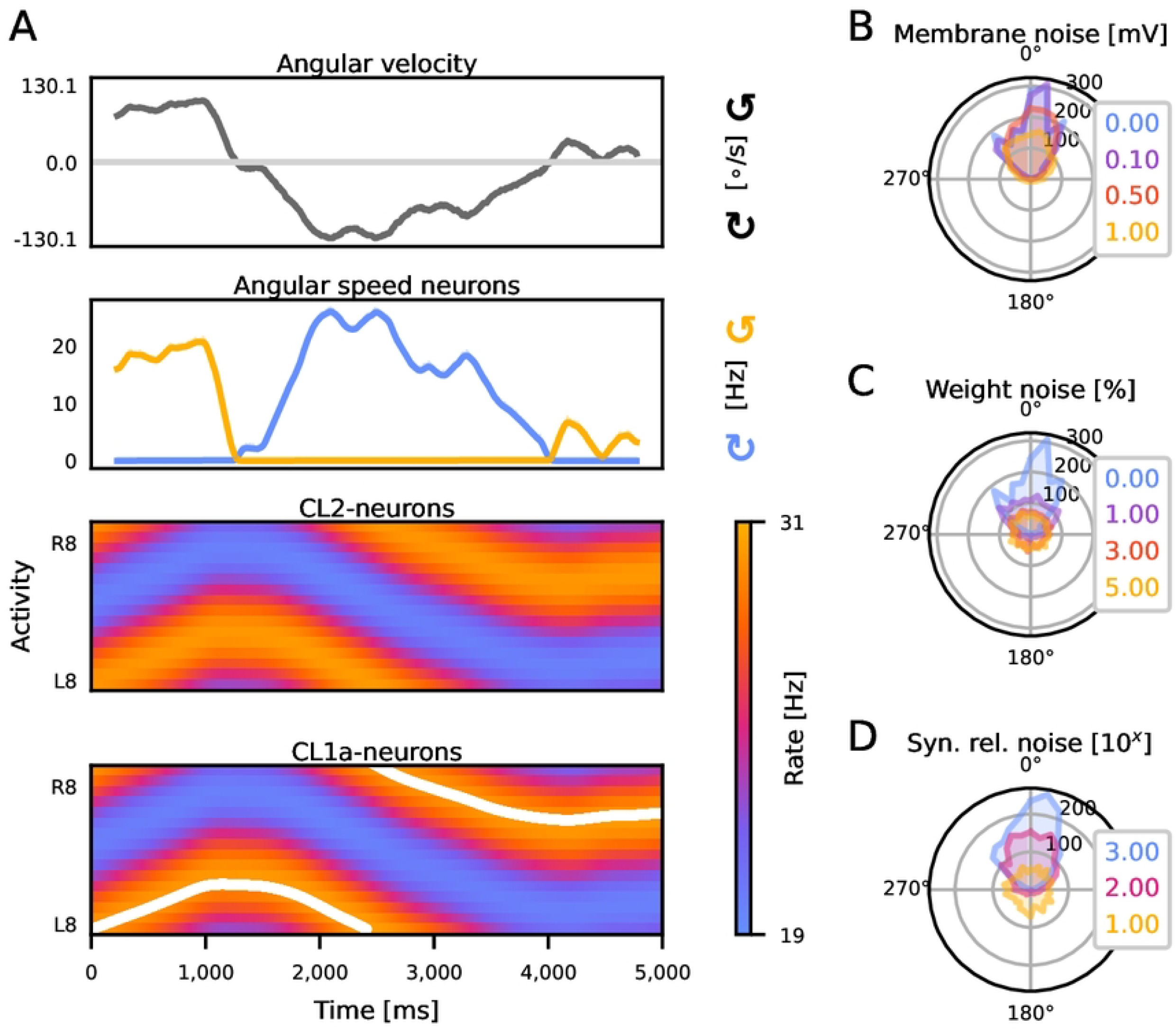
The circuit integrates angular velocity signals to update the heading representation and is resilient to noise. (A) Activity of all neurons of the heading circuit during a noise-free trial of heading- and angular velocity integration. CL1a- and CL2-neurons are organised according to their position in the PB, revealing a single activity bump along the PB. The white line indicates the position of the ideal CL1a compass bump corresponding to the ground truth heading direction computed via Equation 7. (B-D) Accuracy of integration under increasing levels of noise. The angular deviation between the heading encoded by CL1a-neurons and the true heading is depicted (histograms from 1000 trials).

To gauge the robustness of the CL1a+Cl2 modulated circuit’s integration capability, we subjected it to perturbations in three model parameters: membrane potentials, synaptic release probabilities, and synaptic weights. The circuit exhibited graceful degradation [57] in integration accuracy with increasing membrane noise. Membrane noise up to 1 mV and weight noise up to 5 % only marginally increased the number of larger errors in the heading estimate (cf. Figure 5B,C). The model further demonstrated robustness to noise in the post-synaptic channel opening probability, see Figure 5D. Histogram legend shows exponent of pseudocount of noise-generating Beta distribution. Except for very small pseudocounts, i.e., for high probabilities of nonzero noise, the final heading representation pointed in the right direction.

We further investigated the attractiveness of activity states. Figure 6A shows the circuit’s ability to integrate the initial heading encoding and angular velocity inputs under noise-free conditions, resulting in a minimally phase-shifted copy of the initial heading encoding. An illustrative simulation with noise applied to the initial presynaptic inputs producing the initial heading representation is depicted in Figure 6B. Here, the initially noisy heading signal stabilised into an almost ideal sinusoidal activity pattern, signifying the emergence of ring attractor behaviour. Figure 6C displays an example of noise added to synaptic weights, leading to a final CL1a activity state significantly deviating from the initial heading representation or any phase-shifted variant. Given sufficient time, the circuit demonstrated resilience in balancing out noisy initial states caused by perturbations in input rates (cf. Figure 6D). However, the circuit exhibited reduced robustness and accumulated errors when subjected to perturbations in its connectivity induced by noise in synaptic weights (cf. Figure 6E), emphasising the optimality of the optimised circuit connectivity.

**Fig 6.**
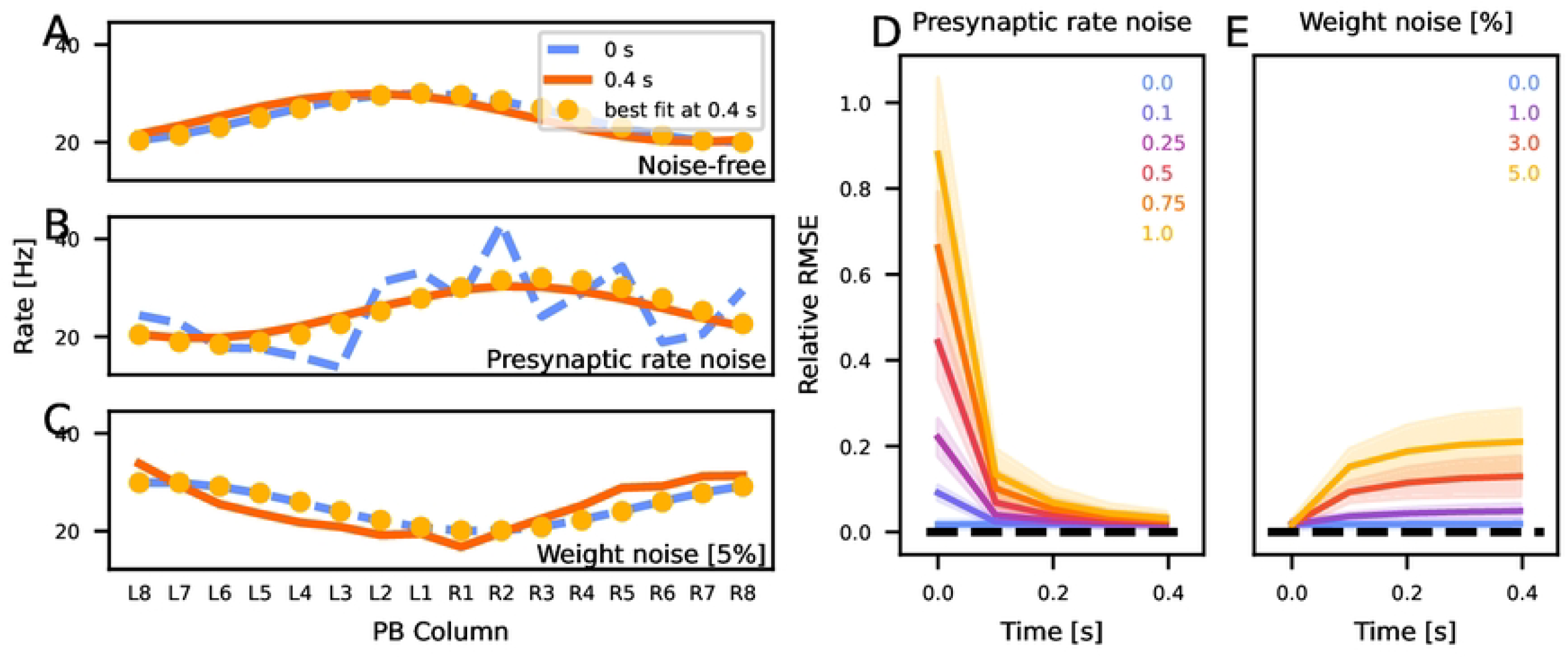
The circuit maintains a sinusoidal activity pattern across the PB. (A-C) show the initial CL1a activity at the beginning of a trial (blue) and the final activity after 0.4 s (yellow). Magenta circles indicate the sine wave best fitting to the final activity. Neurons are arranged according to their position in the PB. Panel A shows an example trial with no added noise, Panel B depicts one with added presynaptic rate noise, and panel C one with added weight noise. (D-E) Development of the relative mean-squared error (RMSE) between the activity of CL1a-neurons at the end of each trial and the best-fitting sine (mean ± SD from 1000 trials). Panel D shows the network converging from noisy initial CL1a activity patterns to less noisy states. Panel E shows that the network activity gets more noisy with added weight noise.

Lastly, we conducted agent simulations to gauge the heading circuit’s efficacy in guiding locomotion in a predetermined goal direction. Figure 4D shows the altered CL1a-to-TB1 connectivity in the goal-directed steering circuit (cf. [41]). In this solution, each TB1-neuron receives excitatory inputs from two adjacent CL1a-neurons, not from one CL1a-neuron from each hemisphere as in the original model (cf. [41] Supplemental Figure S5B). The simulations explored the agent’s ability to maintain a fixed goal direction for 12.5 s. Figures 7A-B show example trials without and with added wind perturbations, respectively. Each trial started with the agent executing a turning manoeuvre to align its heading with the goal direction. Note that the agent lacks the ability to rotate on the spot or to move sideways. Upon being displaced by a wind gust (see Figure 7B), the agent resumed its previous heading. Note that the agent is not equipped with wind sensors and regulates its movements simply by comparing its current heading to the fixed goal heading. The agent exhibited robust performance across diverse probabilities of external translations and rotations, effectively balancing out the effects of perturbations. Figures 7C and D illustrate errors in the heading direction at the end of each simulation. Fig S5A-B illustrate the development of the heading error in time without (A) and with (B) wind-induced perturbations. These results demonstrate the simulated agent’s capability to carry out goal-directed locomotion in dynamic environments, which is important for real-world applications.

**Fig 7.**
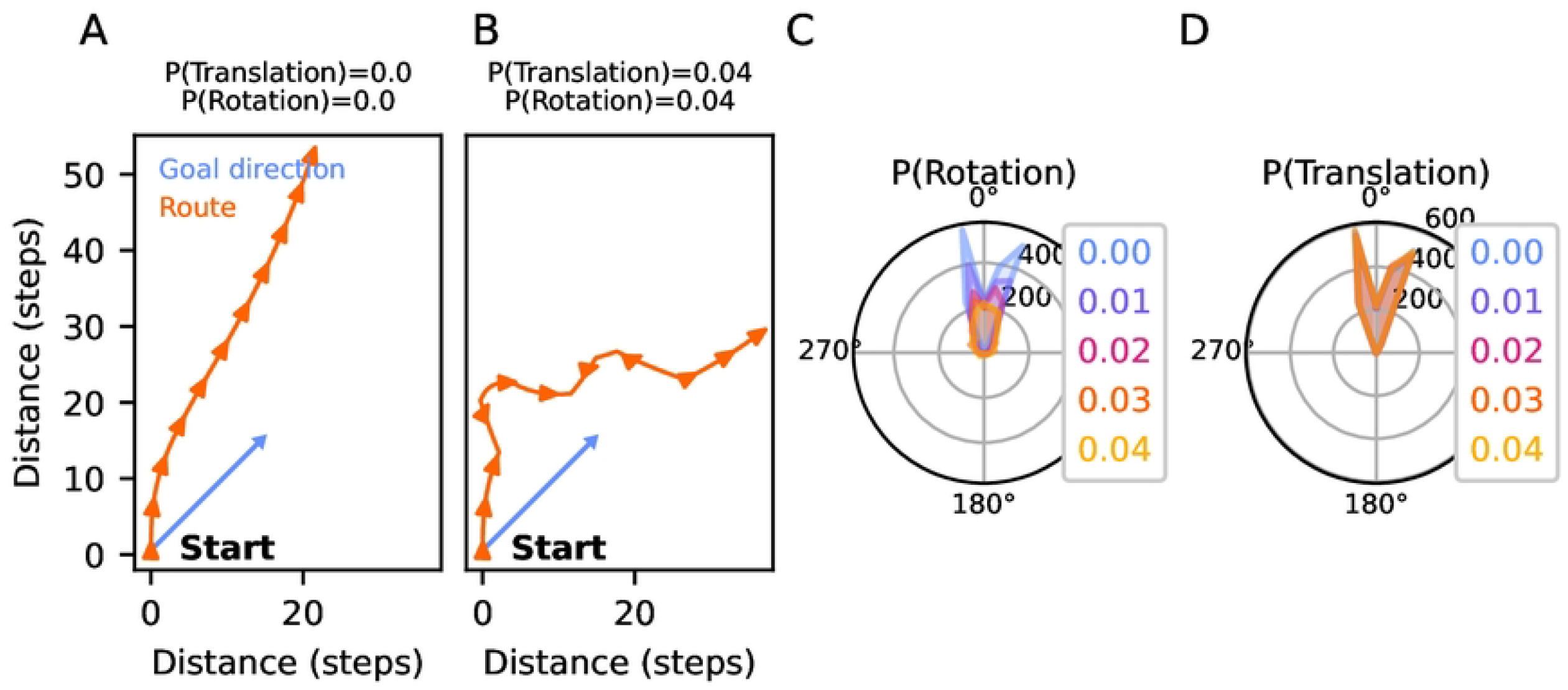
Combined, the heading circuit and the adapted goal-directed steering circuit robustly guide motion in a goal direction, also in the presence of perturbations. (A-B) Example traces illustrating the agent’s motion from a starting location. The agent aligns its initial heading with the goal direction and moves in that direction. Panel A depicts a scenario without external disturbances, while Panel B shows a simulation where gusts of wind cause translation and rotation of the agent. (C-D) Distribution of angular deviations between the goal heading and the agent’s actual heading after 12.5 s (histograms from 1000 trials) under increasing probabilities of translations and rotations of the agent.

The comprehensive evaluations conducted on the proposed circuit consistently demonstrate its robust and reliable performance. While instances of failure emerged under extreme conditions surpassing the network’s noise tolerance, the model exhibits remarkable resilience against minor variations in any parameter or state variable, except for noise added to its connectivity, which emerges as a pivotal network property. These findings underscore the circuit’s potential for stable and accurate functioning in diverse environmental conditions, and highlight the effectiveness of the connectivity solution obtained through optimisation.

## Discussion

Expanding on earlier linear models [36, 43], our study introduces a dynamical synapse firing rate model of angular velocity integration in a locust heading circuit. In contrast to analogous fruit fly models, this novel model exhibits a different compass topography and relies on neuromodulatory, rather than feed-forward, angular velocity inputs.

Our work is situated within the broader context of insect heading circuits, drawing inspiration from models in the fruit fly [2, 33, 58] and other insects [42]. Comparative modeling elucidates adaptations of navigation circuits to species-specific demands, contributing to our broader understanding of adaptive neural circuits. A recent comparative modelling study analysed structural differences between heading circuits in the fruit fly and desert locust [37] but did not account for data suggesting a striking functional difference. Notably, while the fruit fly circuit involves a 2 *×* 360^*°*^ compass mapping along the PB, our model assumes a single 360^*°*^ heading representation. This distinction is implied by physiological data revealing the preferred heading directions of individual CX cells in stationary animals [6, 34, 35, 59]. It remains to be seen whether this fixed 360^*°*^ topography is preserved during active walking or flight. Our study aims to serve as a proof of concept, exploring whether such a topography can produce a stable compass signal.

### Model constraints and properties

The model comprises CL1a- and CL2-neurons along with an abstract class of angular speed neurons. It was constrained by morphological and functional data. Under-constrained connectivity parameters were derived via optimisation.

Heading direction is encoded in the phase of a cosine wave across the PB in our model. The notion of (co)sine waves across neural populations as representations of heading has been a longstanding topic in the literature [60]. Recent advancements, particularly the work by Aceituno and colleagues [61], have underscored their optimality under fairly general conditions, particularly concerning robustness against noise. The optimised circuit successfully updates a heading representation by integrating angular velocity signals and is robust to noise. The fruit fly model by [2] posits asymmetric feed-forward angular velocity inputs to the two halves of the PB that additively modulate firing rates in the heading circuit. Instead of feed-forward inputs, we propose neuromodulatory angular velocity inputs that update the heading signal through multiplicative modulation of firing rates [62] in the locust heading circuit.

In contrast to the fruit fly CX, the locust CX lacks a ring-shaped structure. However, our proposed circuit exhibits key functional ring attractor properties as described in the fruit fly [2, 33, 63]. Our model shares these properties with a previous model of the locust heading circuit featuring a 2 *×* 360^*°*^ compass representation [37]. They include the localisation of input to a single maximum, flexible and cyclic movement of this maximum along the attractor space, and sustained activity in the absence of input. Despite structural deviations from the fruit fly model, the shared ring attractor dynamics suggest convergent solutions to navigation tasks.

Our model of the locust heading circuit could be refined by including recurrent connections from TB1- or TB2-neurons (notice that TB1-neurons are currently not included in the heading circuit itself but appear at its interface with the goal-directed steering circuit). Whether our proposed 360^*°*^ heading representation would still emerge with these recurrent connections remains to be seen. Franconville et al. [12] reported that connections from E-PG-onto P-EN-neurons in the PB are mediated by *Δ*7-neurons rather than being mono-synaptic. Similarly, [58] attribute the same intermediary role to *Δ*7-neurons (referred to as bridge neurons by [9]) in their model of the fruit fly PB. In the locust, TB1- and TB2-neurons may have a similar intermediary function. Notably, TB1a- and TB2-neurons innervate the innermost columns of the PB [54, 64] (cf. Figure 3D). Their role should be investigated in light of new evidence suggesting that the innermost columns in the locust PB consist of two hemi-columns with divergent projection patterns [65].

### Neurobiological plausibility

The effective connectivity of our circuit model is an abstraction of the biological substrate. Neurons in the model can have both excitatory and inhibitory effects on other neurons. Biological implementations could feature additional interneurons, or neurons with opposing effects within the same column of the PB. Electron microscopy studies such as [66] could reassess the model’s connectivity. Likewise, the two angular speed units included in our model, one tuned to clockwise and one tuned to counterclockwise turns, exert up-as well as down-regulating effects on the circuit connectivity through the release of neuromodulators. Such effects could biologically be implemented in tangential neurons innervating the PB (TB3-8-neurons), CBL (TL-neurons) and the lower unit of the NO (NOL, TNL-neurons), most likely through processes more complex than those described in our model. For example, additional interneurons could effectively turn up-into down-regulation. Another explanation could be that the up- or down-regulating effect of the same modulators depends on postsynaptic receptors or signal cascades. An additional possibility is the co-transmission [67, 68] of up- and down-regulating neuromodulators. This assumption contradicts the classical view that each neuron releases a single neurotransmitter, leading to the ‘one neuron, one transmitter’ hypothesis [69], coined as “Dale’s Principle” by [70]. However, many neurons are capable of releasing multiple neurotransmitters [71–74], and this may also be the case in the locust CX. To validate the general concept of tangential neurons acting as angular speed neurons modulating the circuit connectivity, functional studies could assess whether TB-, TL- and TNL-neurons indeed respond to rotation cues and whether they have modulatory effects on columnar neurons.

Furthermore, the recurrent connectivity (see Figure 4A) and the attractor simulations indicate a low-pass filtering of the activity of the CL1a-neuron population. This could be tested in simultaneous neuro-stimulation and multicellular recording experiments, by injecting a noisy activity state into the PB and recording its development in time.

Regarding the employed neuron model, the use of steady-state firing rate neurons is an abstraction and future studies need to verify that the model’s basic principles of heading and angular velocity integration carry over to an implementation with spiking neurons. We have further assumed the same integration and firing dynamics in all neurons, which is likely an oversimplification and might be refined in a more comprehensive CX model. The dynamics of the neurons modelled in our study were constrained by data from homologous neurons in the fruit fly *Drosophila melanogaster* [2]. Obtaining corresponding data from the locust would be crucial to explore potential differences in integration and firing dynamics specific to this species. Testing whether a lead-lag relationship between the activity maxima of E-PG- and P-EN-neurons as reported by [2] also manifests itself in CL1a- and CL2-neurons could be done with multicompartmental models to capture the action potential transmission time along neurites.

### Simulation of goal-directed locomotion

In order to simulate locomotor behaviour, we supplied the heading representation of our model to a goal-directed steering circuit [41]. In this model, a homing vector is encoded in CPU4-neurons and constantly updated. We encoded a fixed goal direction in the CPU4-neuron population to produce steering behaviour as it would be expected during long distance migration, but other cell types are also possible candidates. In the monarch butterfly, goal direction neurons have recently been discovered [75] but were not identified morphologically. However, they could be similar to FC2- and PFL3-neurons in the fruit fly FB (corresponding to CU- and CPU1-neurons in the locust CBU) shown to encode goal directions [5].

In conjunction with the modified goal-directed steering circuit [41], our model can make behavioural predictions. Comparing reaction times of freely moving locusts to shifting visual targets in virtual reality experiments would allow deriving bounds on the functional synaptic delays in our circuit model. Furthermore, such data would allow for the comparison of our model’s feedback control strategy [76] with locust behaviour.

The behavioural simulations conducted in this study are inspired by the initial phase of the proposed three phases of long-range navigational tasks [55]. During this phase, the animal maintains a steady travel direction, guided by global cues. The subsequent short-distance and pinpointing-the-goal phases rely on increasingly specific local cues, underscoring the complexity of successful long-range migration. The evaluations of the simulations we conducted here show that our proposed mechanism for angular velocity integration is robust enough to update the heading signal while other inputs are lacking or are unreliable for a short while. To allow inferences about the circuit’s stability during long-range migration, multimodal cues should be available throughout simulations.

Our current approach involves initializing the activity of the heading circuit based on an initial heading direction and then supplying only angular velocity inputs to update the internal heading signal. To increase the model’s realism, we plan to incorporate sky compass cues into the integration process. This will make the pathway from sky inputs to an internal heading representation with a single compass bump across the PB explicit, inspired by models featuring two compass bumps across the PB [42, 77]. Our understanding of the effective fusion of multimodal cues into a stable heading signal in the desert locust could be furthered by a computational-level analysis (following the framework of [78]) of the heading circuit. This would allow comparisons with similar models and studies of other insects, such as [79] and [80], and exploring the computational principles implemented in the circuit in greater detail. Specifically, adopting an ideal observer model [81] could make the circuit’s objectives under different conditions explicit. Given the potential for conflicting information between different cue modalities in experimental manipulations, simulating experiments under both naturalistic and laboratory conditions would be crucial for a comprehensive evaluation of such a model’s performance.

Identifying fundamental neuronal and computational principles of orientation across different spatial and temporal scales requires future research. Specifically, it should be investigated how the various phases of navigation tasks are integrated, which cues are relevant in each action space [82] and how an animal’s environment, body, and neural system [83] are coupled.

## Supporting information

### Neuron model derivation

The network was modelled based on single-compartment steady-state firing rate neurons abstracted from integrate-and-fire neurons [45]. We outline the abstraction process in the following.

All potentials are measured relative to the membrane reversal potential. Let U be the membrane potential, and c_m_ and g_m_ the specific membrane capacitance and conductance, respectively. For a neuron with synapses indexed by i, g_s,i_ denotes the specific synaptic conductance of synapse i, which we refer to as its weight. P_s,i_ is the synaptic channel open probability, and E_s,i_ is the synaptic reversal potential. The time course of U is then governed by the differential equation (cf. Eq 5.7 in [45]):

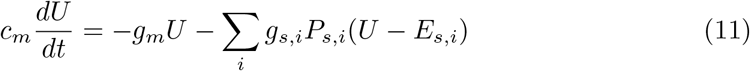

We divide on both sides by g_m_, introduce the membrane time constant 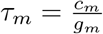 and the relative synaptic weights 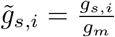 (cf. Eq 5.43 from [45]):

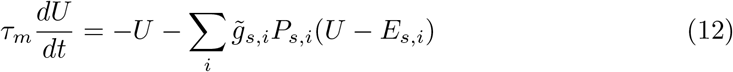

We assume that the membrane time constant *τ*_*m*_ is at least an order of magnitude smaller than the synaptic time constant *τ*_*s*_, which governs the dynamics of *P*_*s,i*_. Estimates for *τ*_*m*_ vary with neuron type. While *τ*_*m*_ = 1.5 ms is common in cortical neurons [45], the membrane time constants used in the fruit fly CX model of [2] are larger by an order of magnitude to capture the observed delays between E-PG and P-EN activity in walking flies [2]. We chose to model these delays by slow synapses, i.e. *τ*_*s*_ *≫ τ*_*m*_. This relationship between the time constants justifies a steady-state approximation to the membrane dynamics. We introduce the steady-state potential

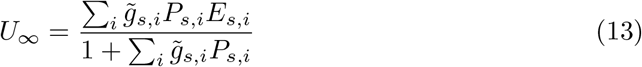

which is reached for constant *P*_*s,i*_ after waiting for a long enough time so that 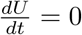.

Using this definition, we can rewrite Equation 12 as

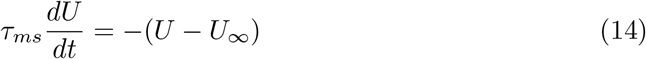

where 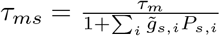 is the membrane time constant corrected for synaptic weights.

Following the derivation in [45], we can then compute the firing rate *r* of an integrate-and-fire neuron whose membrane potential is governed by Equation 14 between spikes via

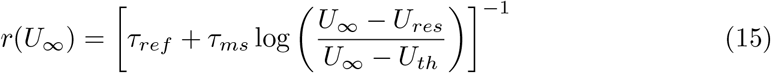

where *τ*_*ref*_ is the refractory period during which the membrane potential is held at the resetting potential *V*_*reset*_ after a spike. *U*_*th*_ is the threshold potential: if *U ≥ U*_*th*_, a spike is triggered. Equation 15 is valid for a deterministic, noise-free neuron. However, thermal membrane potential fluctuations will lead to deviations whose average effect can be well described by a logistic sigmoid:

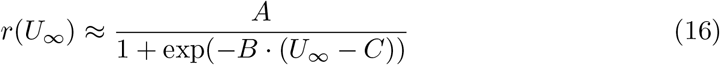

We use the following neuron parameters: *τ*_*m*_ = 1.5 ms, *τ*_*s*_ = 30 ms, *τ*_*ref*_ = 9 ms, *V*_*reset*_ = *−*11 mV, *U*_*th*_ = 15 mV which are typical values for cortical neurons [45]. For thermal fluctuations with a standard deviation of 8 mV, which is the order of magnitude one might expect at room temperature (since the temperature voltage is 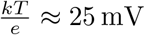) and which has been observed in locust neurons (see e.g. [84], Fig. 2), the best fitting sigmoid parameters are:

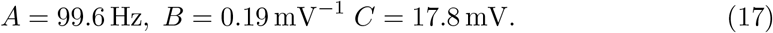

Note that an average rate function with sigmoid shape is obtained for almost any zero mean noise with nonzero standard deviation, so the value of 8 mV is not critical.

An additional bias neuron innervates all columnar neurons. It always fires at a high rate (ca. 100 Hz) to maintain a stable operating point for CL1a- and CL2-neurons (ca. 25 Hz). Other approaches for setting an operating point, such as a change of membrane properties, would be conceivable, too, but are biologically under-constrained.

We describe the time course of the synaptic open probability *P*_*s,i*_ by a single-exponential kernel located at the time of a spike *t*_*spike*_

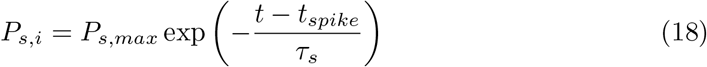

where the synaptic time constant *τ*_*s*_ is typically in the range between 10 to 100 ms. We chose *τ*_*s*_ = 20 ms. Since our network is comprised of rate neurons, individual spike times are not available. We therefore describe the spike timing probability by an inhomogeneous Poisson process parameterised by the (time varying) pre-synaptic rate *r*_*i*_ of synapse *i*. We argue that the Poisson assumption is approximately valid, since *r*_*i*_ does not exceed 50 Hz in available data [36]. Therefore, effects of refractoriness on the regularity of the spike train can be ignored. Following the derivation in [45], the time course of *P*_*s,i*_ can then be described by the differential equation

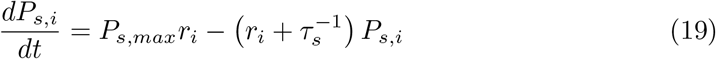

where *P*_*s,max*_ is the maximum synaptic open probability, which we set to 1.

Neurons with a neuromodulatory effect can influence the transmitter release probability *P*_*rel,i,j*_ of a synapse *i* which multiplicatively changes the modulatory weight 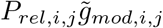. In other words, the total weight 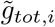 of a modulated synapse *i* is comprised of a constant part 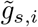 (described above) and a variable part for each modulatory input *j*:

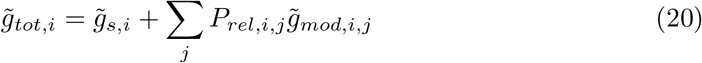

To obtain the steady-state potential in the presence of modulatory synapses, replace 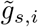 with 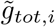 in Equation 1. The time course of *P*_*rel,i,j*_ is governed by a differential equation analogous to *P*_*s,i*_:

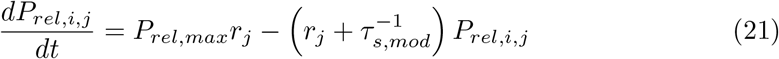

We expect *τ*_*s,mod*_ *> τ*_*s*_ because neuromodulation often involves signal transmission cascades. Therefore, we chose *τ*_*s,mod*_ = *τ*_*s*_ + 10 ms.

Since we did not know a priori whether a synapse would be inhibitory or excitatory, we fixed the synaptic reversal potential of each synapse *i* at *E*_*s,i*_ = 65 mV This is approximately the reversal potential for cations relative to the resting potential [85], and allowed for both positive (excitatory) and negative (inhibitory) synaptic weights 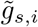 and 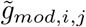. With these signed synaptic weights, the steady-state potential (Equation 13) becomes

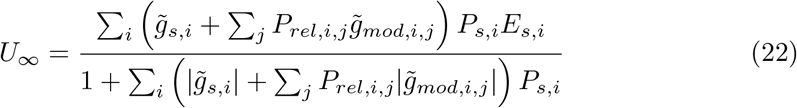

### Functional transmission delays between CL1a- and CL2-neuron populations

Turner-Evans et al. [2] used two fluorescent calcium sensors that radiate at different wavelengths to probe temporal ordering relationships between the activities of P-EN and E-PG neurons of fruit flies walking in the dark. They found that P-EN activity precedes P-EN activity in the EB. In their first experiment, they expressed the fast calcium sensors GCaMP6f [86] in the P-EN neurons, and jRGECO1a [87] in the E-PG population. Fig. 9G of [2] shows a positive angular difference between the compass bumps in P-EN and E-PG that increases with angular velocity, i.e. P-EN is ahead of E-PG. In a second experiment, the calcium sensors were switched between the neuron types (Fig. 9, supplement 2). A smaller, but still largely positive angular difference that increased with angular velocity was found in a small (N=5) number of walking flies.

Here, we tried to use their data for an order-of-magnitude estimation of the effective transmission delay Δ*t* between P-EN and E-PG neurons. We based our estimation procedure on the following assumptions: first, the delay with which a calcium sensor responds to presynaptic spiking activity depends on the sensor, but not on the neuron that is is expressed in. Second, calcium sensor expression does not alter the neuronal dynamics per se. We note that these assumptions may only be approximately correct, see e.g. [86, 87]). Denote the time-dependent angles of the compass bump relative to a reference direction in the P-EN and E-PG population with *ϕ*_*P −EN*_ (*t*) and *ϕ*_*E−P G*_(*t*), respectively. Assume that a fruit fly starts to rotate at orientation *ϕ*_0_ with angular velocity *ω*. Let the unknown transmission delay between E-PG and P-EN be Δ*t*. Then

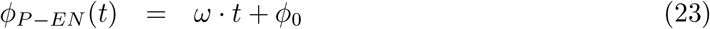

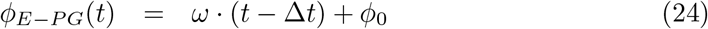

Both of these angles are unobservable. We also assume that the sensor delays Δ*t*_*r*_ and Δ*t*_*g*_ for jRGECO1a and GCaMP6f, respectively, are unknown. The data in [2] are based on the observable flurescence signals. In the first experiment, GCaMP6f is expressed in the P-EN neurons, and jRGECO1a in P-EN. Thus, the time courses of the angles corresponding to the peak amplitudes can be expressed as

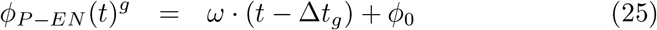

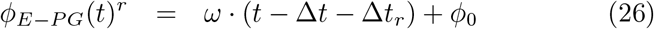

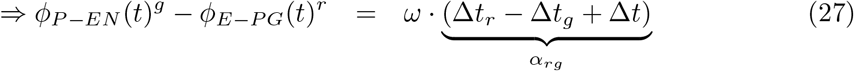

The slope *α*_*rg*_ of the relationship between the angular differences and the (known) angular velocity can therefore be estimated by a linear regression. For the opposite calcium sensor combination, we find

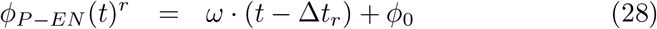

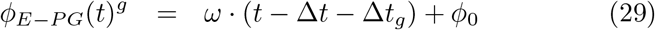

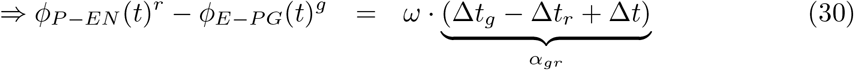

The mean of the slopes therefore is the desired Δ*t*. We estimate the slopes by Bayesian linear regression using the pymc5 package [88], which yields

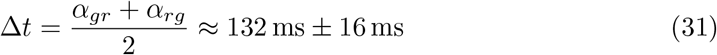

This effective transmission delay is at least an order of magnitude larger than common membrane time constants, which justifies the use of a steady-state neuron model with (slow) dynamical synapses.

---

**Fig S5.**
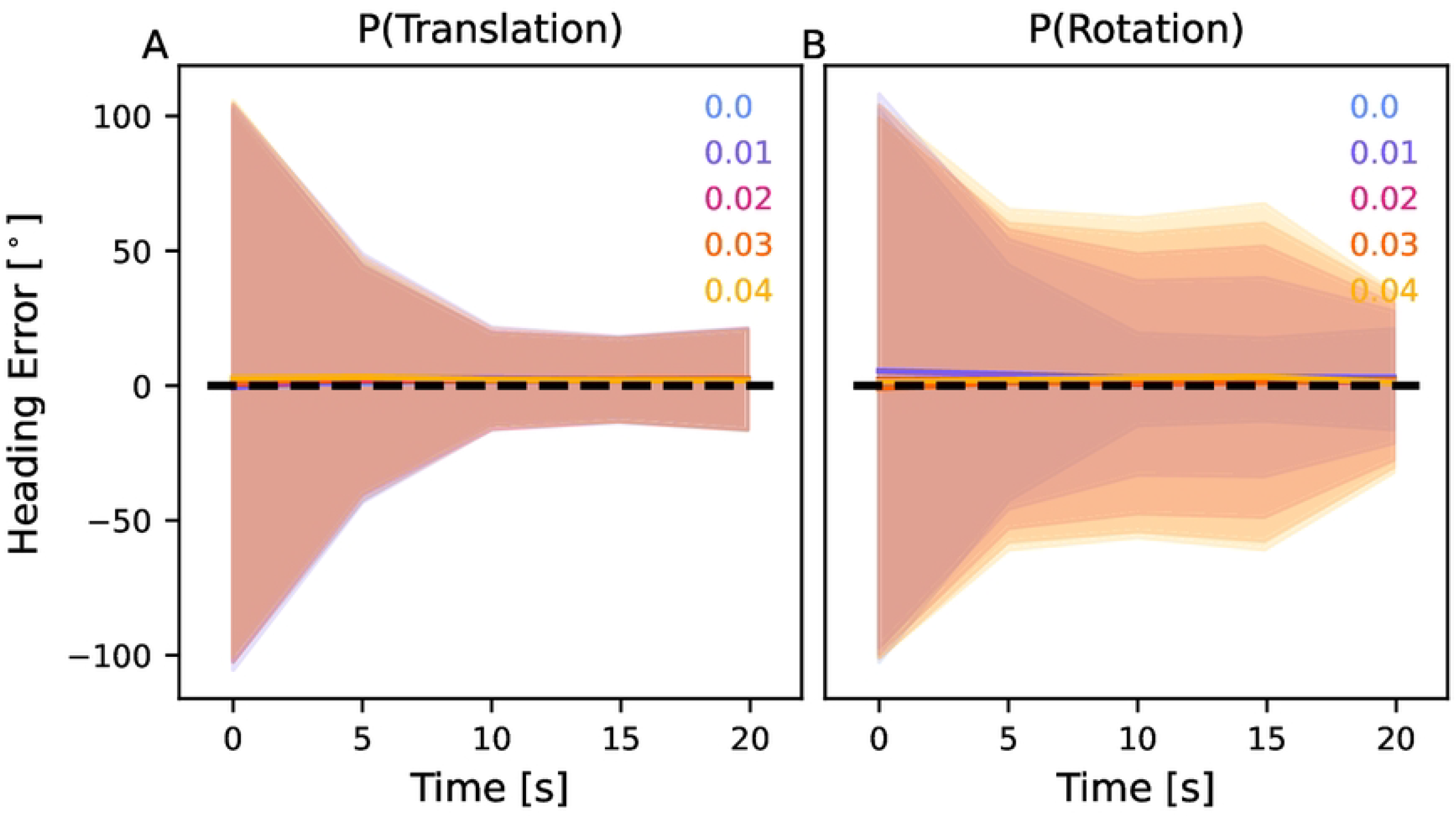
Supplementary Figure to Figure 5. Mean-squared deviation between the agent’s heading estimate and the ground truth heading direction, averaged over 1000 trials lasting 20 s each. Panels A and B demonstrate the impact of the probability of a wind gust that translates or rotates the agent, respectively. Solid lines show the mean angular deviation, shaded areas are one standard deviation. For details, see text.

